# Prediction of drug targets for specific diseases leveraging gene perturbation data: A machine learning approach

**DOI:** 10.1101/2021.12.01.470692

**Authors:** Kai Zhao, Yujia Shi, Hon-Cheong So

## Abstract

Identification of the correct targets is a key element for successful drug development. However, there are limited approaches for predicting drug targets for specific diseases using omics data, and few have leveraged expression profiles from gene perturbations.

We present a novel computational target discovery approach based on machine learning(ML) models. ML models are first trained on drug-induced expression profiles, with outcomes defined as whether the drug treats the studied disease. The goal is to “learn” expression patterns associated with treatment. The fitted ML models were then applied to expression profiles from gene perturbations(over-expression[OE]/knockdown[KD]). We prioritized targets based on predicted probabilities from the ML model, which reflects treatment potential.

The methodology was applied to predict targets for hypertension, diabetes mellitus(DM), rheumatoid arthritis(RA) and schizophrenia(SCZ). We validated our approach by evaluating whether the identified targets may ‘re-discover’ known drug targets from an external database(OpenTargets). We indeed found evidence of significant enrichment across all diseases under study. Further literature search revealed that many candidates were supported by previous studies. For example, we predicted PSMB8 inhibition to be associated with treatment of RA, which was supported by a study showing PSMB8 inhibitors(PR-957) ameliorated experimental RA in mice.

In conclusion, we propose a new ML approach to integrate expression profiles from drugs and gene perturbations and validated the framework. Our approach is flexible and may provide an independent source of information when prioritizing targets.

## Introduction

### Background

Traditionally, drug discovery involves five steps: target identification, target validation, lead identification, lead optimization, and introduction of the new drug to the market [1]. Nevertheless, the speed of new drug development has been slower than anticipated, despite increasing investment [2]. It is estimated that the cost of developing a new drug is ~USD 2.6 billion [3]. One of the main reasons for the enormous cost of drug discovery is the high failure rate.

The success of drug development largely depends on the validity of targets. However, most drugs fail to complete the development process due to a lack of efficacy, and this is often due to the wrong target being pursued [4]. Traditionally, drug targets are often identified from hypothesis-driven pre-clinical models, yet pre-clinical models may not always translate well to clinical applications. For some diseases such as psychiatric disorders, current animal or cell models are still far from capturing the complexity of the human disorder [5]. In addition, some have hypothesized that relying on hypothesis-driven studies alone may have led to ‘filtering’ of findings and publication bias, exacerbating the reliability and reproducibility issues of some research findings [6].

On the other hand, the recent decade has observed a remarkable growth in genomics and other forms of biomedical big data. As increasing amount of data has been made available, computational methods have attracted increasing attention as they offer a fast, cost-effective, and unbiased way to prioritize promising drug targets. Given the limitation of current approaches and the urgent need to develop therapies for diseases, addressing the problem of target identification and drug development from different angles is essential. We believe that computational and experimental approaches can complement each other to improve the efficiency and reliability of identifying valid drug targets. Given the extremely high cost and time investment in drug development, even if the success rate can be increased by a small margin, the savings (in absolute terms) could be substantial.

### Overview of our approach

In this study, we present a novel computational target discovery approach based on machine learning (ML) models to expression profiles induced by genetic perturbation. In our approach, ML models are first trained on drug-induced expression profiles, with outcomes defined as whether the drug can treat the studied disease. The goal is to “learn” expression patterns associated with treatment. The fitted ML models were then applied to expression profiles derived from gene perturbations (i.e., over-expression [OE] or knockdown [KD] of specific genes). We could then prioritize drug targets based on the predicted probabilities from the ML model, which reflects treatment potential.

Intuitively, for example, overexpression (OE) of gene *X* leads to an expression profile ‘similar’ to that of five other drugs known to treat diabetes. Then an agonist targeted at *X* (or other drugs that activate or up-regulate *X* and related pathways) may also be useful for treating diabetes. In this case, we expect the ML model (trained on drugs but applied to gene perturbation data) would output a high predicted probability (of treatment potential) for gene *X*, and it can be prioritized for further studies.

Let us consider an opposite scenario in which over-expression of gene *Y* increases the disease risk or severity. In this case, we may observe a *lower*-than-expected predicted probability of ‘treatment potential’ from the ML model. Gene *Y* can still be considered a potential drug target for further studies, but here we expect *down*-regulation of gene *Y* to be associated with disease treatment.

### Strengths of our approach

Our approach has several potential advantages. Firstly, it provides a general and flexible framework in which any kinds of supervised learning methods can be applied for training. As such, we may leverage the advantages of different, including recently developed, supervised learning algorithms. In addition, our approach is independent of other kinds of evidence usually employed to identify drug targets, for example those used by the OpenTargets platform [7] (e.g., genetic associations, mutation data, expression data, animal models, text mining etc.). The proposed methodology may therefore provide an *independent* source of information when prioritizing targets. Also, our approach does not rely on information of known genes or drug targets for a disease; as such, it may be applicable to a wide range of diseases, including disorders with less well-known pathophysiology and targets. Besides, the lack of reliance on known disease gene/targets may help to discover more novel disease drug targets that are not directly linked to previous ones.

In brief, we first proposed a general framework for identifying drug targets of specific diseases using a machine learning approach, leveraging gene perturbation and drug transcriptome data. Our methodology was applied to several diseases, including type 1 and 2 diabetes mellitus (DM), hypertension (HT), schizophrenia (SCZ), and rheumatoid arthritis (RA). We then validated our new framework by assessing its ability to ‘re-discover’ drug targets based on an external established database (OpenTargets). We also found that many candidate targets are supported by the literature and are functionally relevant.

## Methods

We present a general approach for identifying potential drug targets of a specific disease using state-of-the-art ML methods. As described above, ML models were first trained on drug expression profiles to learn expression patterns associated with treatment of a disease. The trained model was then applied to expression profiles after OE or KD to predict the therapeutic potential of up- or down-regulation of individual genes.

### Datasets

The drug-induced expression profiles and genetically perturbed (OE/KD) expression profiles were downloaded from LINCS (The Library of Integrated Network-Based Cellular Signatures) [8]. For details of the study please refer to [8]. Briefly, to measure the influence of genetic perturbation on expression, each genetic perturbation (OE/KD) was profiled in triplicate 96 hours after application. A single cDNA clone was employed for studies of OE; on the other hand, three distinct shRNAs targeting each gene were profiled for KD experiments. As for expression profiling for drugs, each compound was profiled in triplicate, at 6 or 24 hours following treatment. Gene expression profiling was based on a reduced representation of the transcriptome (1000 ‘landmark’ genes), which has been shown to produce reliable results compared to standard RNA-seq [8].

The original data at multiple levels of pre-processing is available via accession GEO: GSE92742 [8]. In the current study, expression data we was downloaded from the link (https://github.com/dhimmel/lincs) in June 2018, which provides consensus transcriptional signatures for LINCS L1000 perturbations (see https://think-lab.github.io/d/43/#7). Briefly, the input signatures were weighted by its Spearman correlation with other input signatures. For consistency, we kept the genes that appeared in both drug-induced and genetically perturbed expression profiles, so ML models trained on drug expression profiles can be directly employed to make predictions on expression data induced from KD/OE experiments. The final drug expression profile dataset consists of 1158 observations, with expression measured in 7467 genes. The dimensions of the OE and KD datasets were 2413 × 7467 and 4326 × 7467, respectively.

### Training ML models on drug expression data to predict treatment potential

The outcome variable (0/1) is defined as whether the drug is indicated for the disease under study. The drug indications were derived from the Anatomical Therapeutic Chemical (ATC) classification system and the MEDication Indication Resource high precision subset (MEDI-HPS)[17]. We employed our proposed approach to predict drug targets for various diseases covering different systems, including hypertension (HT), type I and type II diabetes mellitus (DM), schizophrenia (SCZ) and rheumatoid arthritis (RA). Indications for HT, DM, and SCZ were extracted from ATC, and indications for RA from MEDI-HPS, because there is no exact category for RA in ATC. We built prediction models for each disease separately, and four ML classification methods were employed for each disease.

#### Model training

The model training procedure largely followed our previous work [9], and we also provide a brief description below. Briefly, we employed four state-of-the-art classification methods, including support vector machine (SVM), gradient boosting machine (GBM), random forest (RF), and logistic regression with the elastic net penalty (EN), to learn the pattern of gene expression profiles associated with treatment of the studied disease[10]-[14]. As the number of drugs known to treat specific diseases is small, there are few observations with positive outcomes. Following our previous study [9], we performed a weighted analysis by increasing weight of the minority class. SVM, RF, and GBM models were implemented using “scikit-learn” in python, while EN was implemented with the R package ‘glmnet’. We employed nested cross-validation (CV) to choose the optimal hyperparameters and evaluate the performance of corresponding models on hold-out datasets. Note that the test-set was only used to evaluate predictive performance and was not involved in hyper-parameter tuning. Please also refer to Supplementary Text for details.

### Model evaluation

In this study, two metrics were used to evaluate the predictive performance of ML models, including area under the receiver operating characteristic curve (ROC-AUC) and area under the precision-recall curve (PR-AUC). PR-AUC may be more instructive in classification performance evaluation when the dataset is imbalanced [15].

### External Validation approach

Validation of drug-disease or drug-target predictions from computational methods has always been a difficult task. As reported by [16], for studies on drug repositioning, a cross-validation approach may overestimate predictive accuracy, as there may be drugs with overlap in the training and testing sets. Also, highly similar drugs may be split into train and test sets; hence, the similarity of training and testing sets may be higher than expected than in practice. There may be similar concerns for disease drug target predictions. If one only evaluates the validity of predictions using performance evaluation metrics (e.g., AUC-ROC) under cross-validation alone, this may lead to over-optimistic results. To avoid this problem, we utilized an independent resource to examine whether our approach can ‘re-discover’ known drug targets for diseases from other data sources. Briefly, we validated our results by evaluating whether the identified targets were enriched for those listed by OpenTargets [7], a platform for systematic drug target identification and prioritization. The platform integrates data from genetics, somatic mutations, expression analysis, drugs, animal models, and the literature through robust pipelines and uses an aggregate score to indicate the association of a target with a disease [7].

Specifically, we applied the models trained on drug expression profiles to OE/KD expression profiles to predict their treatment potentials. Drug targets were downloaded from OpenTargets, with a continuous score (from 0 to 1) indicating the strength of association between the target and disease.

We need to define a cutoff to select relevant genes as ‘valid’ targets for the disease. To avoid arbitrariness in selecting a fixed cutoff, here we defined a cutoff sequence ranging from 1 to 0 with step size 0.2; genes whose association scores lower than the cutoff were filtered away. To test for enrichment, we examined whether the targets from the external database had a higher- or lower-than-expected predicted probability (of treatment potential) from our model, when compared to the non-targets. A two-tailed t-test was used for this comparison. (An alternative approach would be to conduct correlation tests between the target score and predicted probabilities, however the target scores are not normally distributed [with many zeros and ones], which might render such method unreliable.)

## Results

### Model Performance

It should be noted that predictive performance of different ML methods is not the major focus of this study; our main objective is to uncover new disease drug targets and to validate our proposed approach by testing for its ability to ‘rediscover’ known targets.

The average predictive performance of different ML models, measured in AUC-ROC and AUC-PR, is presented in Table S1. In terms of AUC-ROC, SVM performed the best for SCZ, while EN performed the best for DM and RA. GBM slightly outperformed others for HT. In terms of AUC-PR, SVM performed the best in DM and SCZ datasets, but GBM and EN showed the best performance for HT and RA respectively.

### External validation

The results of enrichment test for ‘known’ drug targets from OpenTargets are shown in Tables 1–5. Overall, for drug targets identified from OE data, we observed significant enrichment (with FDR adjusted p-values<0.05) for at least one ML method and score threshold, for all the diseases under study (Tables 1–5).

**Table 1.** Enrichment test of the predicted targets for HT (enrichment for targets listed in OpenTargets)

| Threshold | SVM | RF | GBM | EN |
| --- | --- | --- | --- | --- |
|  | p-value | p-value | p-value | p-value |
| 1 | **4.81E-03** | **3.31E-02** | 9.81E-02 | **2.09E-02** |
| 0.8 | **4.32E-03** | **2.56E-02** | 8.29E-02 | **1.61E-02** |
| 0.6 | **4.41E-04** | **4.94E-03** | *1.76E-02* | **5.68E-03** |
| 0.4 | **8.26E-04** | **4.86E-03** | *1.60E-02* | **1.12E-02** |
| 0.2 | **2.04E-03** | **1.19E-02** | *2.91E-02* | **2.17E-02** |
| 0 | 1.56E-01 | 6.84E-01 | 1.96E-01 | 1.12E-01 |

Enrichment p-values are shown. Four machine learning methods were used to train a model on expression data to predict treatment potential, and the model was fitted to expression profiles after gene perturbation.
FDR adjusted p-values < 0.05 are in bold, while those between 0.05 and 0.1 are in italics.
The first column is the threshold of the ‘relevance’ score (available from OpenTargets) above which we defined a gene as a drug ‘target’.
Enrichment was tested against the targets listed in the OpenTargets database, same below.
SVM: support vector machines; EN: logistic regression with elastic net regularization; RF: random forest; GBM, gradient boosted machines.
For tables 1-5, the predicted targets are based on expression data from over-expression (OE) experiments. Results from KD data are listed in Table S2.

**Table 2.** Enrichment test of the predicted targets for DM

| threshold | SVM | RF | GBM | EN |
| --- | --- | --- | --- | --- |
|  | p-value | p-value | p-value | p-value |
| 1 | *5.78E-02* | *6.53E-02* | **3.28E-05** | **2.32E-03** |
| 0.8 | **2.33E-02** | *2.13E-02* | **4.31E-05** | **1.67E-03** |
| 0.6 | **1.26E-02** | *1.37E-02* | **1.61E-05** | **1.57E-03** |
| 0.4 | **2.32E-02** | *4.32E-02* | **1.72E-03** | **5.22E-03** |
| 0.2 | 6.43E-01 | 3.67E-01 | **1.71E-02** | **3.55E-02** |
| 0 | 3.00E-01 | 6.24E-01 | 8.10E-01 | *8.21E-02* |

**Table 3.** Enrichment test of the predicted targets for RA

| threshold | SVM<br>p-value | RF<br>p-value | GBM<br>p-value | EN<br>p-value |
| --- | --- | --- | --- | --- |
| 1 | 1.18E-01 | **6.23E-04** | **2.01E-02** | 9.22E-01 |
| 0.8 | 1.36E-01 | **3.93E-04** | **8.53E-03** | 9.94E-01 |
| 0.6 | 1.18E-01 | 1.41E-01 | 3.67E-01 | 9.20E-01 |
| 0.4 | 6.44E-01 | **1.14E-02** | **2.25E-02** | 2.69E-01 |
| 0.2 | 3.12E-01 | 8.47E-02 | *4.15E-02* | 8.48E-02 |
| 0 | 3.71E-01 | 7.01E-01 | 1.96E-01 | 2.56E-01 |

**Table 4.** Enrichment test of the predicted targets for SCZ (for targets of SCZ listed in OpenTargets)

| threshold | SVM<br>p-value | RF<br>p-value | GBM<br>p-value | EN<br>p-value |
| --- | --- | --- | --- | --- |
| 1 | 3.32E-01 | 2.84E-01 | 2.57E-01 | 3.56E-01 |
| 0.8 | 4.66E-01 | 2.47E-01 | 2.64E-01 | 2.80E-01 |
| 0.6 | *2.18E-02* | 7.78E-01 | 9.94E-01 | 7.47E-01 |
| 0.4 | *1.91E-02* | 8.62E-01 | 7.97E-01 | 3.84E-01 |
| 0.2 | 7.00E-02 | 8.97E-01 | 5.42E-01 | 7.18E-01 |
| 0 | 7.11E-01 | 1.85E-01 | 9.01E-01 | 3.14E-01 |

**Table 5.**
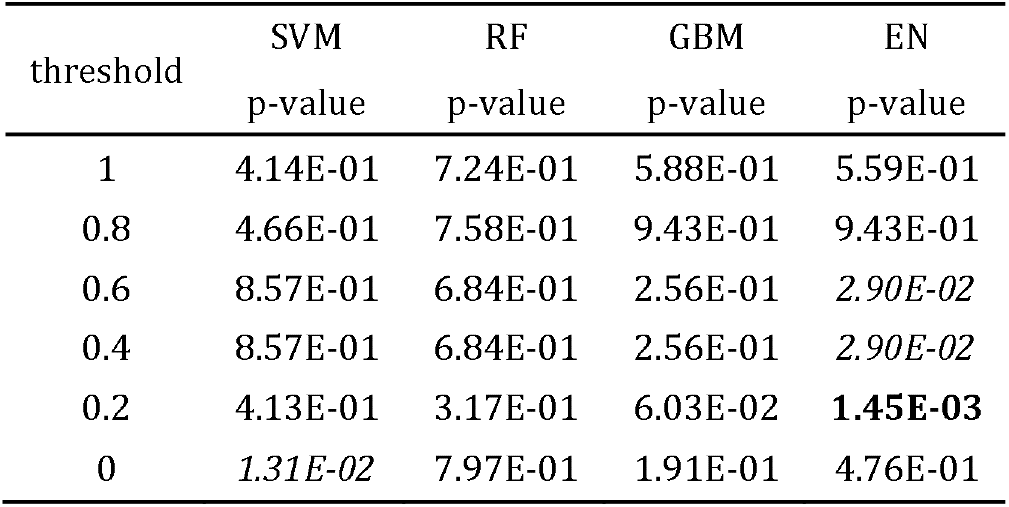
Enrichment test of the predicted targets for SCZ (for targets of bipolar disorder listed in OpenTargets)

| threshold | SVM<br>p-value | RF<br>p-value | GBM<br>p-value | EN<br>p-value |
| --- | --- | --- | --- | --- |
| 1 | 4.14E-01 | 7.24E-01 | 5.88E-01 | 5.59E-01 |
| 0.8 | 4.66E-01 | 7.58E-01 | 9.43E-01 | 9.43E-01 |
| 0.6 | 8.57E-01 | 6.84E-01 | 2.56E-01 | *2.90E-02* |
| 0.4 | 8.57E-01 | 6.84E-01 | 2.56E-01 | *2.90E-02* |
| 0.2 | 4.13E-01 | 3.17E-01 | 6.03E-02 | **1.45E-03** |
| 0 | *1.31E-02* | 7.97E-01 | 1.91E-01 | 4.76E-01 |

On the other hand, apart from a few significant findings for HT, no statistically significant enrichment was observed for targets identified from KD data. The results are shown in supplementary tables (Table S2).

For DM and HT, we observed significant enrichment across multiple thresholds and most of the ML methods with FDR < 0.05, indicating the proposed method indeed ‘re-discovered’ known targets more than expected by chance. For RA, significant enrichment was mainly observed for prediction models based on RF or GBM. For SCZ and BP, which shared anti-psychotics as treatment, the enrichment was not as strong, but suggestive enrichment (FDR < 0.1) was observed especially for SVM in SCZ and EN in BP.

### Literature support of potential targets

In order to validate the functional relevance of our identified potential targets, we conducted a literature search of the 10 targets with the highest and lowest predicted probabilities for each disease (please see Table S3 for a list of these targets), based on targets identified from OE data. As described in the introduction, for targets with high predicted probabilities, we expect that up-regulation of the gene may be associated with therapeutic potential; for targets with lower-than-expected predicted probabilities, we predict that down-regulation of the gene may be associated with treatment.

Selected targets with literature support are discussed below and highlighted in Table 6. Note that our proposed approach does not utilize any prior knowledge of disease-gene associations.

**Table 6.** Literature support of selected drug target candidates

| Potential target | Disease | Direction of expression associated with tx effect (as predicted) | Literature support/Functional relevance |
| --- | --- | --- | --- |
| DRD1 | Schizophrenia | up | Insufficient D1 receptor signaling may be associated with cognitive deficits; D1 agonist has been tested in a clinical trial for cognitive symptoms in SCZ, with moderate improvement in some cognitive tasks observed |
| HIF1AN | Schizophrenia | down | Hypoxia may play a role in SCZ by affecting neurodevelopment; genetic studies showed HIFs may be involved in SCZ |
| ADCY9 | Schizophrenia | up | Involved in glutamate and GABA neurotransmission; *de novo* mutation in the gene may be associated with SCZ |
| SMAD7 | RA | up | *Smad7* expression reduced in synovial tissues of RA patients; mouse models showed that *Smad7* deficiency increased risk to autoimmune arthritis; intra-articular overexpression of *Smad7* relieved experimental arthritis |
| TGFBR2 | RA | up | Linked to resistance of methotrexate treatment and non-responsive patient had reduced expression of the gene in Tregs; hypermethylation (associated with decreased expression) found in RA samples |
| FGFR10P | RA | down | An LD block containing this gene was found in GWAS of RA and other autoimmune conditions. |
| PSMB8 | RA | down | Directly supported by experimental evidence from animal studies: treatment with a PSMB8 inhibitor (PR-957) ameliorated experimental RA in mice. |
| NR0B2 | DM | up | Mutations (associated with reduced activities) in the gene associated with DM; inhibitory effect of metformin on hepatic gluconeogenesis may be mediated through expression of *NR0B2* |
| Fos | DM | up | Insulin induced c-Fos mRNA expression in various cell types including beta-cells; c-Fos upregulation increased beta-cell proliferation, insulin secretion and cellular survival. |
| QPRT | DM | up | Expression of *QPRT* in subcutaneous compartment negatively correlated with HbA1c, fasting glycemia and 120-min glycemia in a clinical study |
| MAGED1 | DM | up | MAGED1-deficient mice showed hyperphagia and reduced motor activity, associated with obesity (shown by two animal studies). *MAGED1* expression was reduced during adipogenesis and loss of MAGED1 led to increased pre-adipocyte proliferation and differentiation *in vitro*. |
| PPP2R1A | DM | up | Encodes a regulatory subunit of PP2A; podocyte-specific loss of PP2A worsened diabetic glomerulopathy and accelerated progression of diabetic kidney disease; interact with IRS1 (Insulin receptor substrate 1), a key mediator of insulin signal transduction implicated in Type 2 DM. |
| TCF7L2 | HT | up | A well-established susceptibility gene for DM found in GWAS (DM and HT are highly comorbid and may shared common pathways); genetic association studies showed associations of SNPs in *TCF7L2* with HT |
| ATP5A1 | HT | up | Reduced expression in HT rats; network analysis showed that actions of a Chinese drug on HT may be mediated through this target |
| FADD | HT | down | Cohort studies reported that a high plasma level of FADD was associated with increased incidence of coronary events and ischemic stroke |

#### Schizophrenia/bipolar disorder

Schizophrenia and bipolar disorder share similar clinical characteristics, and antipsychotics are indicated for both disorders. The two disorders are also highly genetically correlated [17]. We therefore tested for target enrichment for both SCZ and BP based on our model trained on the ATC-SCZ dataset. Our study suggests that overexpression of DRD1 may be associated with treatment effects on SCZ. It was reported that insufficient D1 receptor signaling was associated with cognitive deficits and that working memory deficits may be relieved by treatments that augment D1 receptor stimulation, indicating that drugs acting on this potential target may restore cognitive dysfunction in SCZ [18]. Indeed, DRD1 agonist has been tested in a clinical trial for cognitive enhancement [19]. Moderate improvement was observed on some cognitive tests, including the CogState battery and attention domain of the MATRICS cognitive battery, although no significant improvement was detected for working memory. Other studies also suggested a role of DRD1 in the pathophysiology of SCZ [20]. Taken together, DRD1 may be a potential therapeutic target for SCZ.

HIF1AN, another target identified in our models, has been proven to suppress HIF1A’s transcriptional ability, which can thus affect HIF1A’s ability in regulating hypoxia-inducible genes [21], [22]. Hypoxia may be involved in the pathogenesis of schizophrenia. For example, a methylome-wide association study (MWAS) of SCZ identified many top hits related to hypoxia [23]. *HIF1A* is also proposed as a candidate gene for SCZ, considering the association between *HIF1A* and intrinsic hypoxia occurring in the developing brain that may lead to complex changes in neurodevelopment [24], [25]. HIF1A may enhance vascular growth (hence reducing hypoxia) via controlling the expression of vascular endothelial growth factor (VEGF) [26]. We found that inhibition of *HIF1AN* expression may be associated with therapeutical effect on SCZ, which is in line with the direction of effect from the above studies.

A few other targets may also be associated with SCZ, as supported by other studies. For example, ADCY9 is involved in glutamate and GABA neurotransmission pathways [27], and damaging *de novo* mutations has been identified in the gene [28]. Another potential target *RPA2* showed differential expression in a study of pluripotent stem cell-derived neurons from SCZ patients [29].

#### Rheumatoid arthritis

A number of selected potential targets such as SMAD7, TGFBR2, FGFR10P, and PSMB8 are supported by previous studies. It was reported that *SMAD7* expression was largely reduced in synovial tissues of RA patients, and mouse models also showed that SMAD7 deficiency increased risk to autoimmune arthritis [30]. In addition, it was shown that intra-articular overexpression of *SMAD7* relieved experimental arthritis [31]. These results support our prediction that overexpression of *SMAD7* may improve RA.

Regarding another potential target, TGFBR2, it has been reported that TGFBR2 plays an important role in chondrogenesis [32]. Current results showed that up to 40% of RA patients are resistant to methotrexate, the first-line therapy for RA. Peres et al. reported that drug-resistance of methotrexate was linked to a reduction of CD39 expression due to the impairment in TGF-β signaling, and TGF-β increases CD39 expression on regulatory T cells (Tregs) via the activation of TGFBR2 [33]. The authors also observed that patients non-responsive to methotrexate had reduced expression of *TGFBR2* in Tregs compared to responsive patients. In this connection, overexpression of *TGFBR2* may reverse impairment of TGF-β signaling, which is consistent with our prediction that *TGFBR2* overexpression may be useful for RA treatment. In addition, hypermethylation of TGFBR2 (associated with decreased expression) was found in RA samples [34]. Taken together, the results above indicate that *TGFBR2* expression levels might be linked to RA disease activity.

Additionally, FGFR10P was identified as a possible target for RA by our study. An LD block on chromosome 6 (6q27) which contains the genes *CCR6* and *FGFR1OP* was observed to be associated with increased risks for several autoimmune diseases, such as RA, Crohn’s disease and vitiligo [35]-[40].

Moreover, our model predicted that inhibition of PSMB8 may induce treatment effects on RA. This is directly supported by experimental evidence from animal studies. It was observed that treatment with a PSMB8 inhibitor (PR-957) can ameliorate experimental RA in mice [41]. The drug led to decrease in cellular infiltration, cytokine production and autoantibodies in the RA mouse model. In a similar vein, several studies also reported that PSMB8 inhibitors reversed autoreactive immune responses and showed therapeutic effects in animal models of autoimmune encephalomyelitis, colitis and Hashimoto’s thyroiditis [42]-[44]. Regarding evidence from human genetics studies, a SNP in the *PSMB8* (*LMP7*) gene was also found to be associated with juvenile RA [45].

#### Diabetes mellitus

Previous studies also support several potential targets identified by our approach, and our study suggested that overexpression of these targets may be associated with treatment effects on DM. First, Mayumi et al. found that mutations of *NR0B2*, also known as *SHP*, was associated with type 2 DM in a Japanese sample. It was reported that the mutant proteins show significantly reduced activities [46]. Another study showed that the inhibitory effect of metformin (one of the most commonly used drugs for DM) on hepatic gluconeogenesis may be mediated through expression of NR0B2 [47].

We also identified Fos as a potential target for DM (with OE favoring treatment). It was found that insulin could induce *c-Fos* mRNA expression in neurons [48], fibroblasts [49] and pancreatic beta-cells [50]. Another study showed that *c-Fos* upregulation increased beta-cell proliferation, insulin secretion and cellular survival, mediated by the activation of Nkx6.1. On the other hand, *c-Fos* knockdown inhibits Nkx6.1-mediated beta-cell proliferation and reduces insulin secretion [51].

QPRT (quinolinate phosphoribosyltransferase) was also found to be a potential target for DM. It is an enzyme involved in the kynurenine pathway, which may be involved in diabetes pathogenesis [52]. In a recent clinical study, the expression of *QPRT* in the subcutaneous compartment was negatively correlated with HbA1c, fasting glycemia, and 120-min glycemia[52]. This is consistent with our finding that up-regulation of this target may be associated with therapeutic potential.

We also revealed MAGED1 as a potential drug target, and we predicted that up-regulation of the gene may be associated with treatment of DM. One study concluded that MAGED1-deficient mice showed hyperphagia and reduced motor activity, which led to the development of obesity [53]. Another subsequent animal study [54] observed a similar phenomenon that MAGED1-deficient mice showed late-onset obesity, owing to reduced energy expenditure and physical activities. The study also found that *MAGED1* expression was reduced during adipogenesis and loss of MAGED1 led to increased pre-adipocyte proliferation and differentiation *in vitro*. MAGED1 also reduced the stability and transcriptional activity of PPAR-gamma, which is the target for thiazolidinediones (a class of anti-diabetic medication).

Several other identified targets were also shown in previous studies to be associated with DM. For example, *GADD45A* was suggested as a diabetes-associated gene, which might be involved in both diabetic cardiomyopathy and DM-induced baroreflex dysfunction [55]. Regarding another potential target, *TSPAN8*, a SNP in this gene was associated with insulin release and sensitivity in a genetic association study [56]. Another target of interest was PPP2R1A, which encodes a constant regulatory subunit of protein phosphatase 2 (PP2A). It was found that podocyte-specific loss of PP2A worsened diabetic glomerulopathy and accelerated the progression of diabetic kidney disease [57]. In addition, PPP2R1A was discovered to interact with IRS1 (Insulin receptor substrate 1), a key mediator of insulin signal transduction implicated in Type 2 DM [58].

#### Hypertension

Our models identified TCF7L2 as a potential target for hypertension. *TCF7L2* is a well-established susceptibility gene for type 2 DM [59], and given the high comorbidity rate and possibly shared pathophysiology between DM and hypertension [60], further studies into this target may be warranted. In addition, a study of the Thai elderly population suggested that the SNP rs290487 in *TCF7L2* may contribute to risks of hypertension regardless of Type 2 DM [61]. Another cohort study concluded that both a parental history of diabetes and the *TCF7L2* at-risk variant were associated with a higher incidence of hypertension, after controlling for other cardiometabolic risk factors [62].

Considering another potential target, ATP5A1, a pharmacological network analysis of Compound Uncaria Hypotensive Tablet (a Chinese medication for hypertension) revealed that the therapeutic effect of this drug may be associated with actions on ATP synthetases including ATP5A1 [63]. Another study revealed that the expression of *ATP5A1* was significantly decreased in spontaneously hypertensive rats compared with controls [64]. These results may further support our finding that overexpression of *ATP5A1* may be associated with therapeutical effects.

Another target of interest is FADD, which is also a marker of apoptosis and apoptosis may be implicated in atherosclerosis [65]. Cohort studies reported that a high plasma level of FADD was associated with increased incidence of coronary events and ischemic stroke [66], [67]. Considering the strong associations between hypertension, stroke, and coronary heart diseases [68], the findings from these studies are supportive of the role of FADD and our prediction that inhibition of FADD expression may be associated with treatment of hypertension (or its complications).

Interestingly, TSPAN8, which was suggested by our approach as a DM drug target, was also identified as a potential target for hypertension. As argued above, given the comorbidities and possibly shared metabolic pathways [60], TSPAN8 may also be an interesting candidate.

Some other potential targets, such as DUSP6 and HOXB13, are also supported by the literature. Zoe et al. [69] found that genes from the DUSP family may contribute to hypertensive heart disease; specifically, for *DUSP6*, its expression was up-regulated in spontaneously hypertensive rats compared to controls. The above results were consistent with our finding that the inhibition of *DUSP6* may have a protective treatment effect. For HOXB13, it was reported that knockdown of *HOXB13* can reduce the cytotoxicity caused by various oxidative stress inducers [70], [71], and an increasing number of studies suggest that oxidative stress has a key role in the pathogenesis of hypertension [72].

## Discussion

### Overview

In this study, we presented a novel ML-based computational approach to identify promising drug targets. To our knowledge, this work is the first to employ ML methods to leverage both drug-induced and genetically perturbed expression data to discover potential drug targets for specific diseases.

Our approach is general as it can incorporate any supervised learning algorithms. To validate our method, we examined whether it may ‘re-discover’ known targets based on other sources of data. Indeed, we observed that top genes from our models were enriched for targets from the OpenTargets platform. Encouragingly, a number of targets highlighted by our proposed method were also supported by the literature.

### Relevant works

We highlight a few relevant works on drug target prediction here. Kandoi et al. reviewed machine learning and system biology applications in distinguishing drug targets from non-targets [73]. Several studies explored biological properties of known drug targets by ML methods to predict druggability of proteins [74]-[78]. For example, Kumari et al. proposed a sequence-based prediction model and leveraged information like amino acid composition and amino acid property group composition to predict whether a new target may be druggable. They also performed a comprehensive comparison of several ML methods [77]. In another study, eight key properties of human drug targets were extracted and learned by SVM to discover new targets [74], [78] . A similar study extracted simple physicochemical properties from known drug targets to predict targets against non-targets [78].

Regarding network-based approaches, Costa et al. [79] leveraged interaction network topological features together with tissue expression and subcellular localization data to predict druggable genes. In another work, Li et al. employed topological features of protein-protein interaction network to identify potential drug targets [76]. Emig et al. presented an integrated network-based method to predict drug targets based on disease gene expression profiles and an interaction network, and some novel drug targets for scleroderma and cancer were reported [80].

However, our study is different from the previous studies in several aspects. One of the most important differences is that the focus of most of the above studies (except [80]) is to predict *in general* whether a protein may serve as a drug target (i.e., distinguishing targets from non-targets). They did not address the problem of predicting whether a protein is a target *for a specific disease*, such as diabetes or schizophrenia. As discussed above, network-based methods are useful and has been proposed for uncovering disease drug targets. However, they are relatively dependent on similarity between entities and known drug targets, hence may be less capable of discovering novel targets. Also, network-based approaches usually require good knowledge of gene-gene (or protein-protein) and disease-gene interactions. It may not be easy to define such interactions accurately and different sources may suggest different patterns of interactions. The edges may therefore need to be defined arbitrarily.

There are relatively few studies that employed gene perturbation data to predict drug targets, but a recent study [81] has leveraged such data to identify tentative targets. The authors proposed pairwise learning and joint learning methods constructed on chemically and genetically perturbed gene expression profiles to predict targets of different chemicals [81]. They also constructed a drug-protein-disease network for drug repurposing. However, the methodologies and objectives of our study and ref [81] are different. We proposed ML methods to assess how the expression profiles from gene perturbations are related to those of drugs. Ref [81] mainly employed Pearson correlation and linear models to assess the similarity between transcriptomic changes from gene perturbation and those from drugs. An advantage of our approach is that by employing ML methods (e.g., SVM, random forests, boosted trees), we may accommodate complex non-linear relationships and interactions between features. Study [81] used transcriptomic data from gene perturbations mainly to predict drug-protein interactions; prediction of disease-specific drug targets was performed in a separate analysis using networks (which requires knowledge of the known therapeutic targets of studied diseases). As discussed above, network-based methods have their own limitations. We proposed an alternative new approach, which integrates transcriptomic data with ML approaches in a unified framework to predict drug targets *for specific diseases*.

One of our previous works has employed an ML approach for drug repositioning, leveraging drug expression data [18]. However, the objectives are different from the current study, in which we aim to uncover novel drug targets. In practice, drug repositioning may not always be feasible (for example, due to side-effects of existing drugs), and there are also important hurdles to drug repositioning efforts, such as patent considerations, regulatory barriers and organizational hurdles in industry, as reviewed by Pushpakom et al. [82]. As a result, revealing new targets remains a very important goal in drug development and pharmaceutical research. Besides, unlike our previous work, here we have covered diseases other than psychiatric disorders. Also, gene perturbation data has not been used in the previous study.

### Strengths and limitations

As described earlier, there are important strengths of our approach. Our approach is general and highly flexible, can incorporate any supervised ML methods, is independent of other sources of evidence commonly employed to identify drug targets, and does not rely on knowledge on known disease genes/targets. However, there are also several limitations. One limitation is that our ML prediction model building datasets are highly imbalanced, as only a small number of drugs are usually indicated for each disease. In order to address this issue, we increased the class weight of the minority group. There are other strategies to address issues, such as SMOTE (Synthetic Minority Oversampling Technique) [83], but whether strategies like SMOTE can address this issue in high-dimensional settings is still unclear and will be a topic for further investigations. Another aspect is that we observed significant enrichment for the identified targets primarily in the OE datasets but not in KD datasets. One hypothesis is that some off-target effects may interfere with the expression profiles in KD experiments, leading to greater difficulties in finding relevant drug targets [84]. How to overcome or reduce the influence of off-target effects remain an area for further studies. Here we have employed enrichment tests to examine ‘re-discovery’ of known drug targets from other sources of data, and showed that many targets may be clinically/biologically relevant based on the literature. Nevertheless, further experimental and clinical studies are required to confirm our findings. Also, further works are required to elucidate the mechanisms underlying the drugs that may act on the identified targets.

## Conclusion

This study presented a general computational framework to prioritize drug targets for various diseases. Under the framework, different kinds of ML methods can be utilized. We applied four ML methods to identify potential drug targets of four disorders. External validation showed that the top candidates are enriched for targets selected by independent lines of evidence from a large external database (OpenTargets). We also found that previous studies provided support to a number of targets identified by our approach.

Finding promising targets for diseases is crucial to drug development. However, it is impractical to perform indepth experimental studies on every possible target for each disease. Computational methods offer a cheap, fast, and systematic high-throughput approach to guide the prioritization of targets. We hope our presented framework will provide an additional way to prioritize drug targets, which may benefit future drug development.

## Author Contributions

Conception and design: HCS (lead) and KZ. Study supervision: HCS. Funding acquisition: HCS. Methodology: HCS, KZ. Data analysis: KZ. Data interpretation: HCS, YS, KZ. Preparation of first draft of manuscript: KZ and HCS, with input from YS.

## Acknowledgements

This work was supported partially by a National Natural Science Foundation China (NSFC) grant (81971706) and the Lo Kwee Seong Biomedical Research Fund from The Chinese University of Hong Kong and the KIZ-CUHK Joint Laboratory of Bioresources and Molecular Research of Common Diseases, Kunming Institute of Zoology and The Chinese University of Hong Kong, China.

