## Supplementary material for "Prediction of drug targets for specific diseases leveraging gene perturbation data: A machine learning approach": Supp Text and Supp Tables 1-3: supp text OE KD paper.docx

**Supplementary Text**

**Hyperparameter tuning and nested cross-validation**

SVM, RF, and GBM models were implemented using “scikit-learn” in python [1], and we performed a two-step hyperparameter tuning with gridsearchCV provided in the package [2]. For SVM [3], we chose the radial basis function (RBF) as the kernel for our model, and the hyper-parameters C and gamma were chosen from (-5, 15) and (-20, 2) in log-2 space, respectively. For RF [4], we fixed the number of trees to 1000, and selected the maximum number of features (max_features) for each splitting and minimum number of samples for each leaf (min_samples_leaf) from {800, 1000, 1500, 2000, 3000, 5000} and {1, 3, 5, 10, 30, 50, 80} respectively. For GBM [5], learning rate was chosen from {0.005, 0.01, 0.015, 0.02, 0.03, 0.05}, the number of boosting iterations from the sequence from 100 to 1001 with step size 50, the maximum depth of each estimator from {2, 3, 5, 10} and maximum number of features from {10, 30, 50, 100, 500, 1000}. The subsampling proportion was fixed to 1. Finally, we implemented EN [6] using the R package “glmnet”, with hyperparameter α ranging from 0 to 1 with step size 0.1 and λ following the default setting. Some refinements of the parameters grid of the above models were carried out after analyzing model fitting.

In one of our previous studies, we used nested cross-validation (CV) [7] to choose optimal hyperparameters and evaluate the performance of corresponding models on hold-out datasets. Specifically, in a nested CV, we chose the hyperparameters based on the performance of the corresponding models on the validation set in the inner CV; in the outer CV, a hold-out test set was used to evaluate the predictive performance of the optimal model chosen in the inner CV. Here we adopted the same strategy since simple CV tends to overestimate model performance, and it is more informative to present more precise accuracy of the optimal model [8]. The splitting of datasets is the same across all models by setting the same random seed, which guarantees a fair comparison of the performances of our prediction models.

**List of Supplementary Tables**

Table S1	Average predictive performance of different machine learning methods across four datasets

Table S2	Enrichment test results of the predicted targets from knockdown (KD) data

Table S3	List of identified targets (the 10 targets with the highest and lowest predicted probabilities of treatment potential are shown; four sub-tables showing targets for each disease)
