## Supplementary material for "Prediction of drug targets for specific diseases leveraging gene perturbation data: A machine learning approach": Supp Text and Supp Tables 1-3: Table S1 pred performance table .docx

Table S1 Average predictive performance of different machine learning methods across four datasets

|  | ATC DM | ATC HT | MEDI-HPS  RA | ATC SCZ |
| --- | --- | --- | --- | --- |
| Average AUC-ROC | | | | |
| SVM | 0.6232 | 0.5433 | 0.5709 | 0.7582 |
| RF | 0.6024 | 0.5488 | 0.5706 | 0.7377 |
| GBM | 0.5404 | 0.5516 | 0.5244 | 0.7474 |
| EN | 0.6485 | 0.5506 | 0.5788 | 0.7496 |
| Average AUC-PR | | | | |
| SVM | 0.0834 | 0.0804 | 0.0649 | 0.2402 |
| RF | 0.0616 | 0.0884 | 0.0471 | 0.2113 |
| GBM | 0.0578 | 0.0937 | 0.0485 | 0.2106 |
| EN | 0.0338 | 0.0792 | 0.0706 | 0.2362 |

The figure for the best performance of learning algorithms in each dataset for different evaluation metrics is in bold.

ROC-AUC: area under the curve (AUC) of the receiver operating characteristic (ROC) curve; PR-AUC: area under the curve (AUC) of the precision-recall (PR) curve.

SVM: support vector machines; EN: logistic regression with elastic net regularization; RF: random forest; GBM, gradient boosted machines.

MEDI-HPS: MEDication Indication - High Precision Subset; ATC: Anatomical Therapeutic Chemical classification.

DM stands for diabetes mellitus, HT for hypertension, SCZ for schizophrenia, and RA for rheumatoid arthritis.
